# Defining distinct RNA-protein interactomes of SARS-CoV-2 genomic and subgenomic RNAs

**DOI:** 10.1101/2023.05.15.540806

**Authors:** Isabella T. Whitworth, Rachel A. Knoener, Maritza Puray-Chavez, Peter Halfmann, Sofia Romero, M’bark Baddouh, Mark Scalf, Yoshihiro Kawaoka, Sebla B. Kutluay, Lloyd M. Smith, Nathan M. Sherer

## Abstract

Host RNA binding proteins recognize viral RNA and play key roles in virus replication and antiviral defense mechanisms. SARS-CoV-2 generates a series of tiered subgenomic RNAs (sgRNAs), each encoding distinct viral protein(s) that regulate different aspects of viral replication. Here, for the first time, we demonstrate the successful isolation of SARS-CoV-2 genomic RNA and three distinct sgRNAs (N, S, and ORF8) from a single population of infected cells and characterize their protein interactomes. Over 500 protein interactors (including 260 previously unknown) were identified as associated with one or more target RNA at either of two time points. These included protein interactors unique to a single RNA pool and others present in multiple pools, highlighting our ability to discriminate between distinct viral RNA interactomes despite high sequence similarity. The interactomes indicated viral associations with cell response pathways including regulation of cytoplasmic ribonucleoprotein granules and posttranscriptional gene silencing. We validated the significance of five protein interactors predicted to exhibit antiviral activity (APOBEC3F, TRIM71, PPP1CC, LIN28B, and MSI2) using siRNA knockdowns, with each knockdown yielding increases in viral production. This study describes new technology for studying SARS-CoV-2 and reveals a wealth of new viral RNA-associated host factors of potential functional significance to infection.

## INTRODUCTION

Severe acute respiratory syndrome-related coronavirus 2 (SARS-CoV-2) is the causative agent of coronavirus disease 2019 (COVID-19) which has been associated with over 6.8 million deaths since its emergence in 2019.^1,2^ Like all viruses, SARS-CoV-2 encodes a limited number of proteins and relies on interactions with host factors for efficient replication. Interactions between SARS-CoV-2-encoded RNAs and host proteins mediate nearly all aspects of the viral life cycle including viral mRNA synthesis, translation, genome replication, and infectious virion assembly. Host proteins also play central roles in defense against viral replication through the activation of antiviral gene programs or directing the mutation or degradation of viral RNAs.^3,4^ Characterizing the protein interactomes of viral RNAs is thus crucial to understanding fundamental aspects of virus replication as well as mechanisms of cell intrinsic and innate antiviral immunity.

SARS-CoV-2 is an enveloped, positive-sense (+), single-stranded RNA beta coronavirus with a 30 kilobase (kb) genome. Upon infection of host cells, the SARS-CoV-2 genomic RNA (gRNA) serves as an mRNA template decoded to produce the viral nonstructural proteins (NSPs) essential for replication.^5,6^ In total, 16 NSPs are translated from two large open reading frames (ORFs) with the majority of NSPs interacting to form viral replication and transcription complexes (RTCs). The gRNA also serves as the template for synthesis of negative-sense (-) RNA intermediates, driven by the RTC. The (+) RNAs are then synthesized using (-) RNAs as templates. In total, 11 (-) RNA species are produced to yield 11 (+) RNAs. These 11 viral RNA species include new copies of the full-length (+) gRNA and 10 (+) subgenomic RNAs (sgRNAs).^7,8^ The sgRNAs are generated through a mechanism termed discontinuous transcription, resulting in a 3’ coterminal nested set of 10 canonical sgRNAs that share a 5’ leader sequence (Figure 1A).^7,9,10^ Four of the viral sgRNAs code for viral structural proteins: Spike (S) protein, Nucleocapsid (N) protein, Matrix (M) protein, and Envelope (E) protein. These proteins package the gRNA and form new virions that are released by the cell. The remaining sgRNAs code for viral accessory proteins that vary in number and function between coronavirus species and are largely believed to contribute to modulation of host cell responses.^10^

**Figure 1.**
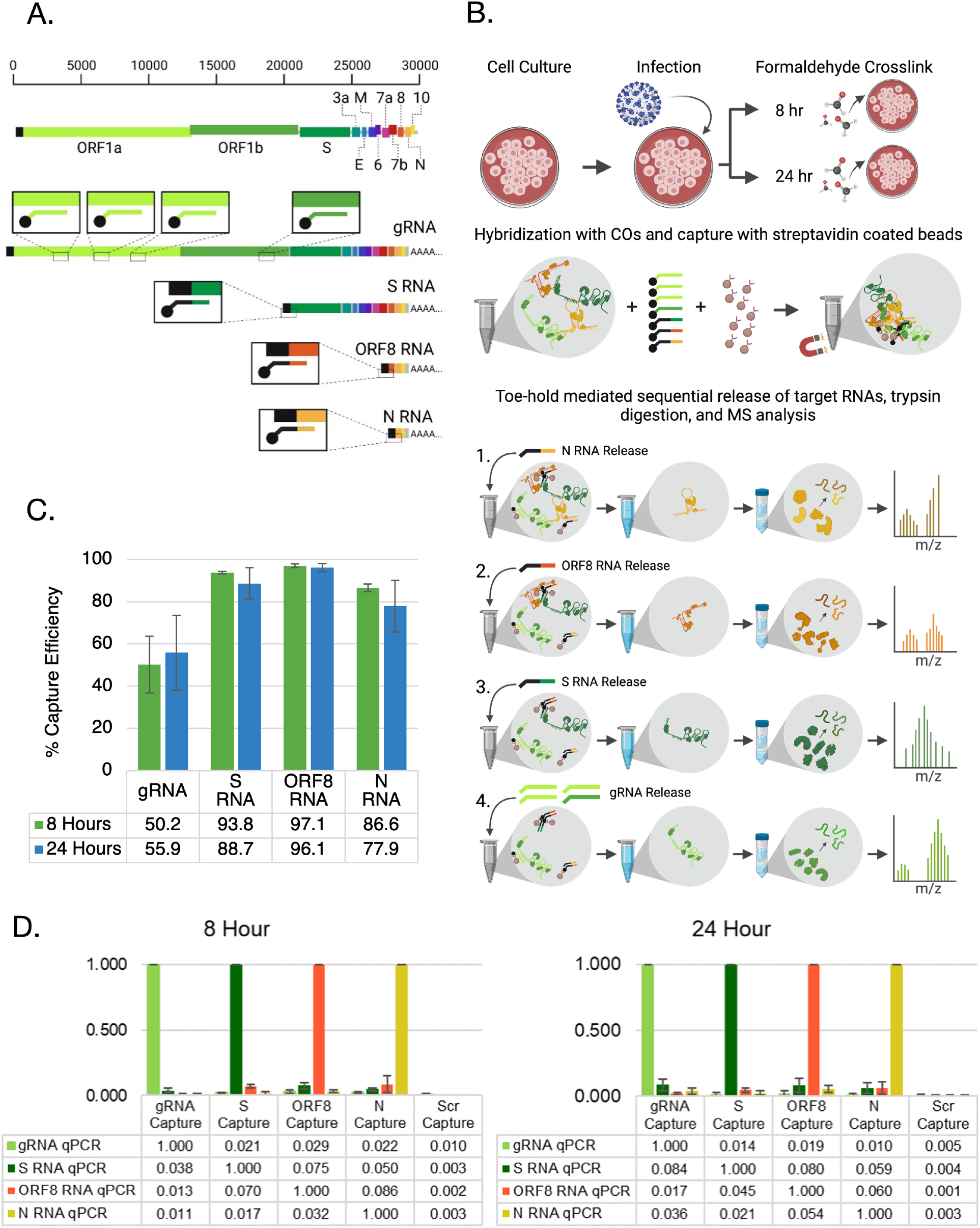
HyPR-MS_SG_ for purification of SARS-CoV-2 sgRNA interactomes. (A) Capture oligonucleotides (COs) were designed to bind to specific regions of the SARS-CoV-2 gRNA or three individual sgRNAs to allow for simultaneous, specific capture of all four RNAs individually from a single infected cell lysate. (B) Description of the HyPR-MS_SG_ procedure. (C) Capture efficiency was determined by measuring the extent of the depletion of the target RNA from the pre-capture lysate using RT-qPCR. (D) Capture specificity was measured by comparing the ratio of the targeted RNA to non-targeted RNAs in each capture using RT-qPCR. Error bars in (C) and (D) represent standard deviations of the mean from the four biological replicates.

The specific protein interactome of each (+) viral gRNA and sgRNA is predicted to be highly dependent on the individual RNA’s identity and function. Additionally, the protein interactome of each of these RNAs is likely to significantly vary over the course of infection. For example, host cells are first infected and sgRNA production is established after formation of the RTC and generation of the (-) sgRNA intermediates. However, later in the course of infection, viral sgRNAs are translated to produce structural and immunomodulatory proteins, copies of the gRNA are produced, and virion production begins. For SARS-CoV-2 in cell culture systems, infectious viral particles have been reported as early as 6-hours post infection (hpi), but peak replication levels are achieved between 24- and 48-hpi.^11,12^

To better understand how the virus interfaces with host cell machinery, prior studies have examined the net effects of SARS-CoV-2 infection on the host proteome or transcriptome and studied viral RNA-protein interactions by isolating gRNAs individually or combined with all other viral sgRNA interactomes mixed.^13–15^ However, to date no study has been able to separate and characterize individual sgRNA pools and associated proteins. Because of the important and distinct role that sgRNAs play in the SARS-CoV-2 life cycle, a comprehensive understanding of virus-host interactions necessitates resolution at the level of individual sgRNAs and at multiple stages of the virus life cycle. To address this gap, herein we describe a strategy for identification of specific RNA-protein complexes to identification of specific sgRNA-protein complexes, <u>hy</u>bridization purification of <u>p</u>rotein-<u>R</u>NA complexes followed by <u>m</u>ass <u>s</u>pectrometry for coronavirus <u>sg</u>RNAs (HyPR-MS_SG_). For validation, we distinguish the specific protein interactomes of SARS-CoV-2 gRNA and three independent sgRNA pools (S, N, and ORF8) at 8- and 24-hours post infection, with these two time points selected to capture both early viral infection when RNA synthesis is just beginning and viral replication is being established, as well as a later stage where protein synthesis and virion packaging is thought to occur. We identified 509 proteins in total associated with these viral RNAs, with around half of these proteins previously characterized as SARS-CoV-2 RNA binders including expected viral proteins such as N and components of the viral replicase. In total, 260 proteins were identified that had not been previously characterized as interacting with SARS-CoV-2. Gene ontology analysis showed an overrepresentation of proteins linked to translation initiation, RNA splicing, protein kinase binding, posttranscriptional gene silencing, and regulation of ribonucleoprotein granules. siRNA-mediated depletion of select protein interactors with putative antiviral functions resulted in significant increases in viral RNA levels and consequently viral proteins. This study provides an unprecedented resolution to characterize individual SARS-CoV-2 sgRNA-protein interactomes and reveals multiple new functionally significant virus-host interactions.

## MATERIALS AND METHODS

### Cell lines, viruses, and infection

All cell lines were maintained in a humidified incubator at 37°C at 5% CO_2_ unless otherwise indicated. The Huh7+AT cells used for HyPR-MS_sg_ experiments were Huh7 cells (a gift from Dr. Daniel Loeb, University of Wisconsin) modified to express the SARS-CoV-2 receptor ACE2 and entry co-factor TMPRSS2 after transduction with ACE2- and TMPRSS2-encoding lentiviral vectors (gifts of Caroline Goujon, Montpelier University, France).^16^ A derivative of Vero E6 cells (ATCC-CRL-1586) engineered to stably express human TMPRSS2 (a gift of Dr. Sean Whelan) was used for propagation of SARS-CoV-2 strain 2019-nCoV/USA-WA1/2020 (obtained from the Centers for Disease Control and Prevention and a gift from Dr. Natalie Thornburg), and SARS-CoV-2 mNeonGreen (SARS-CoV-2-NG) reporter viruses.^17,18^ Huh7+AT cells were cultured in Dulbecco’s modified Eagle essential minimal medium nutrient mixture with Ham’s F-12 medium (DMEMF12; Gibco) supplemented to a final concentration of 10% with heat-inactivated fetal bovine serum and 1% penicillin-streptomycin-L-glutamine (PSG) solution. Calu-3 (ATCC-HTB-55), Vero (ATCC-CCL81) and Vero E6 (ATCC-CRL-1586) cells were cultured in Dulbecco’s modified Eagle medium supplemented with 10% fetal bovine serum. After infection, cells were washed with media to remove unbound virions and media was replaced. For HyPR-MS studies, Huh7+AT cells were seeded at density of 10^7^ cells in a 10 cm dish. Twenty-four hours later, cells were infected with SARS-CoV-2 at an MOI 1.0 i.u./cell. For virus propagation, cells were infected at an MOI of 0.01 i.u./cell and cell culture supernatants were collected after visible cytotoxicity was reached, at ∼3 days post-infection. Virus stock titers were determined on Vero E6 cells or Vero E6-TMPRSS2 by plaque assays and sequenced to confirm identity to the reference sequence. Calu-3 cells transfected with siRNAs were infected at a MOI of 2 i.u./cell for an hour at 37°C, after which the initial inoculum was removed, washed once with 1X phosphate buffered saline (PBS), and replaced with cell culture medium.

### Cell lysis, hybridization, capture, and release of genomic and subgenomic RNA-protein complexes

At 8-hpi and 24-hpi, cells were washed three times with cold PBS, with proteins cross-linked by resuspending the cells in 0.25% formaldehyde and incubating at room temperature for 10 min. Cells were then scraped off the dish and centrifuged to pellet the cells. The pellet was washed three times with cold PBS and then quenched for ten minutes in 100 mM Tris-HCl, incubated at room temperature. Cells were washed twice in 1xPBS, pelleted by centrifugation, and frozen at −80°C. The lysis, hybridization, capture, and release protocols are adapted from the HyPR-MS protocol in Henke *et al*., summarized here to include any changes. Cells were lysed in and sonicated as previously described.^19^ Capture oligonucleotides (COs) targeting gRNA, S sgRNA, N sgRNA, and ORF8 sgRNA and negative control scrambled sequence oligonucleotides (SOs) were added to the cell lysates (Supplementary Table 1). The samples were then incubated at 37°C for three hours with gentle nutation. Sera-Mag Streptavidin Coated Magnetic Speedbeads (Fisher Scientific) were washed twice at RT and added to the lysate. The samples were rocked for one hour before bead collection and lysate removal. Beads were then washed and resuspended in release buffer with release oligonucleotides (ROs) (Supplementary Table 1). The beads were gently rocked at RT for 30 minutes to release RNA-protein complexes, then the supernatant was isolated and stored at 4 °C for eventual RT-qPCR and mass spectrometric analyses. This process was first done with N ROs then repeated 4 subsequent times with ORF8 sgRNA, S sgRNA, gRNA, and scrambled RO oligonucleotides, respectively resulting in capture of all targets from the same lysate.

### RNA extraction, reverse transcription, and qPCR analysis

A small fraction (2%) of each release sample was incubated overnight at 37°C with 1 mg/mL Proteinase K (Sigma), 4 mM CaCl_2_, and 0.2% LiDS to remove proteins. The RNA was then extracted from each sample using TRI Reagent (Sigma) per the manufacturer’s protocol and precipitated in 75% ethanol with 2μL of GlycoBlue coprecipitant (Thermo Fisher Scientific) at −20°C for at least 2 hr. The RNA was pelleted by centrifugation at 20,800 g at 4°C for 15 min, washed with 75% ethanol, centrifuged at 20,800 g at 20°C for 15 min, then resuspended in 15μL of nuclease free water (Invitrogen). 10μL of the purified RNA was used for reverse transcription (High-Capacity cDNA Reverse Transcription Kit, Applied Biosystems) per the manufacturer’s protocol. The 20μL reverse transcription product was diluted with 60μL of Invitrogen nuclease free water and analyzed using sequence-specific qPCR primers and probes (Integrated DNA Technologies) and Light Cycler 480 Probes Master Mix (Roche) for relative quantitation of RNA on a CFX96 Touch real-time PCR detection system (Bio-Rad) (Supplementary Table 1).

### Protein purification, tryptic digestion, and mass spectrometry of peptides

The remainder of each capture sample was processed using an adapted version of eFASP for purification of proteins, as previously described.^20^ For removal of salts from the sample, an OMIX C18 solid-phase extraction pipette tip (Agilent) was used according to manufacturer’s instructions. The samples were then dried in a SpeedVac and reconstituted in 95:5 H_2_O: acetonitrile (ACN), 0.1% formic acid. The samples were analyzed using an HPLC-ESI-MS/MS system consisting of a high-performance liquid chromatography (nanoAcquity, Waters) set in line with an electrospray ionization (ESI) Orbitrap mass spectrometer (QE-HF orbitrap, ThermoFisher Scientific). A 100μm id × 365μm od fused silica capillary micro-column packed with 20cm of 1.7μm diameter, 130 Å pore size, C18 beads (Waters BEH), and an emitter tip pulled to approximately 1μm using a laser puller (Sutter Instruments) was used for HPLC separation of peptides. Peptides were loaded on-column with 2% acetonitrile in 0.1% formic acid at a flowrate of 400nL/min for 30 min. Peptides were then eluted at a flowrate of 300 nL/min over 150 min with a gradient from 2% to 30% acetonitrile, in 0.1% formic acid. Full-mass profile scans were performed in the orbitrap between 375 and 1500 m/z at a resolution of 120,000, followed by MS/MS collision induced dissociation scans of the 10 highest intensity parent ions at 30% relative collision energy for the first replicate and 35% relative collision energy for the second replicate and 15,000 resolution, with a mass range starting at 100 m/z. Dynamic exclusion was enabled with a repeat count of one over a duration of 30 s.

### Mass Spectrometry Data Analysis

Mass spectral files were analyzed with the free and open-source search software program MetaMorpheus and the reviewed Swiss-Prot human XML (canonical) database.^21^ Samples were searched allowing for a fragment ion mass tolerance of 20 ppm and cysteine carbamidomethylation (static) and methionine oxidation (variable). A 1% FDR for both peptides and proteins was applied. Up to two missed cleavages per peptide were allowed for protein identification and quantitation. To determine the differential interactomes of the SARS CoV-2 RNAs, pairwise comparisons of each SARS-CoV-2 RNA compared to the scrambled control were statistically analyzed with the student’s t-test using Perseus software.^22^ Proteins that met the threshold of 20, 20, 10, and 5% FDR for gRNA, S RNA, ORF8 RNA and N RNA, respectively, or a student’s t-test statistic in the pairwise comparisons were then evaluated for Gene Ontology enrichment and protein-protein interactors using STRING.^23^ For each protein, the intensities across all capture samples were normalized to the N protein intensity.

### siRNA transfections, infections, and downstream analyses

1x10^5^ Calu3 cells were reverse transfected in 24-well plates with 5pmoles of siRNAs using RNAimax (Thermo-Fisher) using manufacturer’s instructions (Supplementary Table 1). Media was replenished following overnight incubation with siRNAs and infections were initiated 48-hours post-transfection as detailed above. Cells were collected at 24-, 48-, 72-, and 96-hpi for RNA extraction, immunoblotting or fixed by 4% paraformaldehyde for FACS analysis. RNA was extracted by TRIzol (Invitrogen) using manufacturer’s instructions. RNA was reverse transcribed using High-Capacity cDNA Reverse Transcription Kit (Applied Biosystems). Resulting cDNA was subjected to qRT-PCR on a ViiA Real-Time PCR System (Thermo Fisher) using gene-specific primers diluted in POWERUP SYBR Green master mix (Thermo-Fisher). For immunoblotting cells were lysed in 1X radioimmunoprecipitation buffer (50 mM Tris pH7.4, 1% NP-40, 0.25% Na-deoxycholate, 0.1% SDS, 150 mM NaCl, 1mM EDTA). Lysates were separated on 4-12% Bis-Tris polyacrylamide gels and transferred to nitrocellulose membranes. Immunoblotting was performed as previously described.^24^ Primary antibodies used were Rabbit polyclonal anti-SARS2 nucleocapsid, Sino Biologicals/40588-T62, and Mouse monoclonal anti-actin: SantaCruz/SC-8432. Membranes were probed with fluorophore-conjugated secondary antibodies (LI-COR) and scanned using an LI-COR Odyssey system. IN and CA levels in virions were quantified using Image Studio software (LI-COR).

## RESULTS

### Purification of SARS CoV-2 genomic and subgenomic RNA-protein complexes

We previously described HyPR-MS as a strategy to elucidate the protein interactomes of individual pools of mRNAs, lncRNAs, viral RNAs and viral RNA splice variants.^25–28^ Here, we adapted the approach to allow for isolation of specific sgRNA pools and associated protein interactomes from a single population of SARS-CoV-2 infected Huh7+AT cells (with this modified technique henceforth referred to as HyPR-MS_SG_). Huh7+AT cells were selected for this analysis due to their capacity to support high levels of viral replication using the 2019-nCoV/USA-WA1/2020 virus strain. Four SARS-CoV-2 target RNAs were selected for HyPR-MS_SG_ based on their crucial roles in the viral life cycle: gRNA, S sgRNA, ORF8 sgRNA, and N sgRNA. The S sgRNA codes for the Spike protein which covers the surface of virions and interacts with host cell proteins to mediate viral entry into cells.^29^ The N sgRNA codes for the highly abundant RNA-binding Nucleocapsid protein that coats viral RNA, regulating transcription and gRNA packaging into virions.^30^ The ORF8 sgRNA codes for the ORF8 viral accessory protein which is found in a highly variable region of the genome and may represent an important virulence factor.^31^

Because the sgRNAs of SARS CoV-2 are formed through discontinuous transcription of the SARS-CoV-2 gRNA, each sgRNA contains an identical 5’ leader sequence positioned upstream of a protein coding region followed by a 3’-polyA tail (Figure 1A). Each of the resulting sgRNA shares 3’ sequence homology with the gRNA and all other SARS-CoV-2 sgRNAs of greater length, except at the junction between the leader sequence and the body of the RNA (henceforth called the leader junction). ^7,9^ Therefore, to achieve efficient and specific sgRNA isolation, a single HyPR-MS_sg_ capture oligonucleotide (CO) specific to each sgRNA target was designed to span each unique leader junction. Unlike the sgRNAs, the large unique region of gRNA (approximately 20,000nt) allowed for the design and combined use of multiple specific COs, so that four independent COs were designed to hybridize to the gRNA’s unique region (Figure 1A). Additionally, a scrambled CO was designed with a similar melting temperature and GC content to that of the target COs, to serve as a control for nonspecific protein binding. After RNA capture, RT-qPCR assays specific for S, N, and ORF8 sgRNAs and the gRNA were used to confirm RNA capture specificity and efficiency (Supplementary Table 1). For the four biological replicates, capture efficiencies were >75% for each sgRNA and >50% for gRNA (Figure 1C), with all captures enriched at least 10-fold relative to off-target RNAs, and with only low levels of SARS-CoV-2 RNAs detected in the scramble CO captures (Figure 1D). These data demonstrated that each targeted CO or set of COs captured its corresponding SARS-CoV-2 RNA pool with the specificity needed for mass spectrometry and comparative protein interactome analysis. Accordingly, the remaining capture samples were processed for analysis using protein purification and trypsin digestion, with the resulting peptides analyzed by bottom-up mass spectrometry to define each SARS-CoV-2 RNA-protein interactome.

### Characterization of the protein interactomes for specific SARS CoV-2 RNAs

Two technical replicates using two different collision energies were analyzed by mass spectrometry for each of four biological replicates. Individual protein intensities were summed for each sample and then normalized to viral N protein intensity, considering that N is a known binder of SARS CoV-2 RNA and thus served as proxy measure for total viral RNA captured. Scrambled oligonucleotide captures were normalized to overall total protein intensity.

Using a student’s t-test and permutation-based false discovery rate (FDR) statistical analysis, we compared protein intensities from all four biological replicates for each individual viral RNA capture to the scrambled oligonucleotide capture controls. Two tiers were established to categorize statistically significant changes to protein abundance. The first tier was comprised of proteins that passed a 20, 20, 10, and 5% FDR cutoff set for gRNA, S RNA, ORF8 RNA and N RNA captures, respectively. The second tier was determined using a sliding scale that considered both fold difference and p-value, and thus was comprised of proteins with a t-test statistic of >2.78 (Figure 2A, Supplementary Table 2). Based on these criteria, the total protein interactomes (tiers 1 and 2 combined) for SARS-CoV-2 genomic RNA and the S, ORF8, and N sgRNAs were defined as being comprised of 58, 65, 65, and 93 proteins, respectively, at 8-hpi; and 35, 15, 331, and 290 proteins, respectively, at 24-hpi (Figure 2A, Supplementary Table 2). To further assess selectivity, we determined which proteins derived from each individual viral RNA interactome were unique vs. also enriched in one or more additional SARS-CoV-2 RNA interactome at either time point (Figure 2B). Interestingly, at 8-hpi, approximately half of the proteins in the gRNA and S sgRNA interactomes were identified as significant only in a single interactome but dropping at 24-hpi, while ORF8 and N sgRNAs exhibited relatively large abundances of both unique (>25% of total) and shared proteins (>50% of total) at the later (24-hpi) time point.

**Figure 2.**
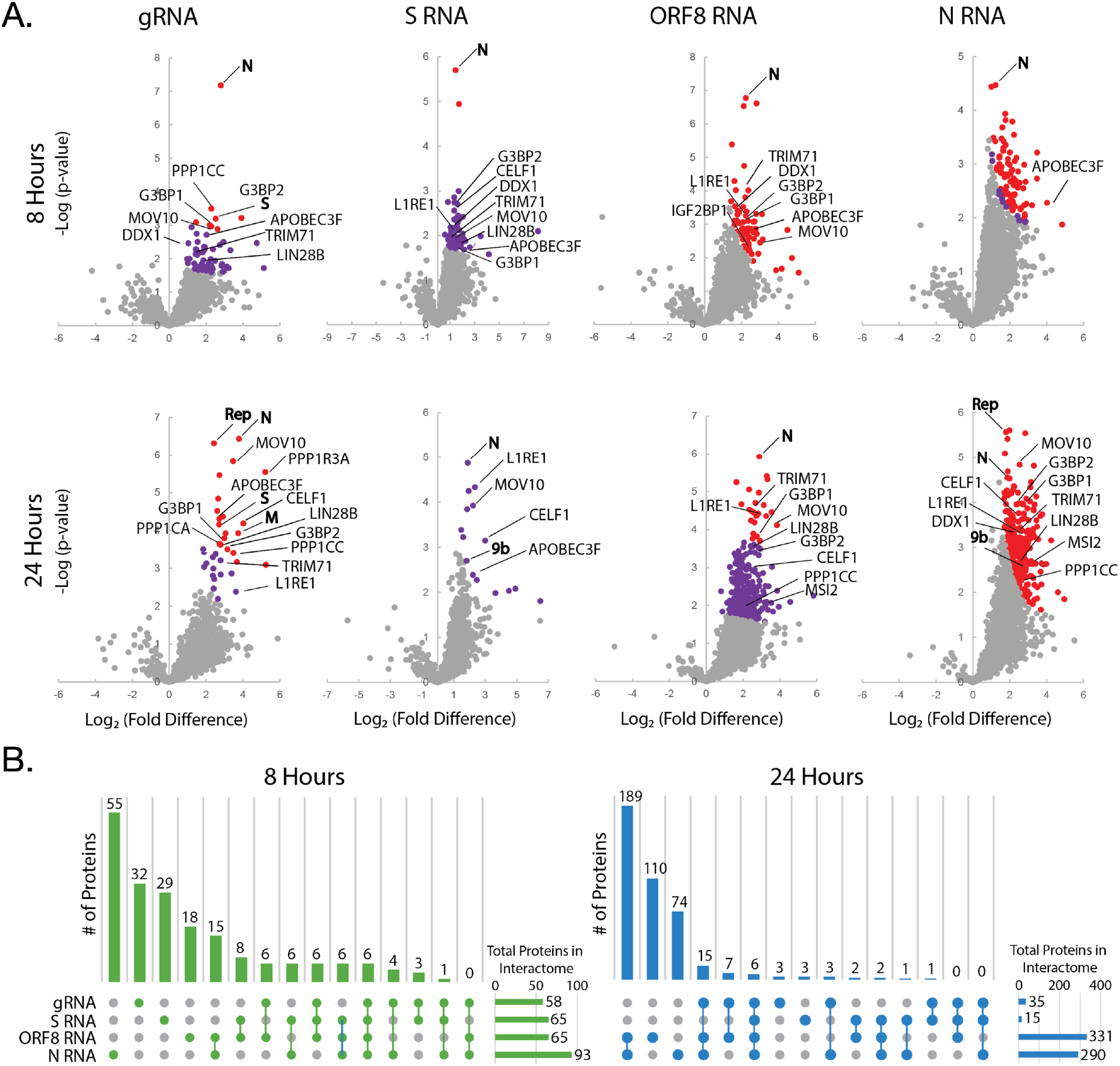
Determination of SARS-CoV-2 RNA protein interactomes. (A) Volcano plots displaying the distribution of all proteins for each SARS-CoV-2 RNA capture with relative protein abundance (log_2_ target RNA capture/scrambled control capture) plotted against significance level (-log_10_ p-value), with significantly enriched proteins indicated by colored dots. Red dots indicate proteins that passed a 5% FDR cutoff using a student’s t-test (tier 1) and purple dots indicate proteins that passed the less rigorous fold-change and p-value cutoff metrics (tier 2). (B) Graph displaying the number of proteins found in each individual or subset of interactomes for each of the two time points.

We also expected viral proteins to be identified in each specific RNA interactome. N protein, which we would expect to be bound to all viral RNAs, was a top hit in all interactomes (Figure 2A). The tier 1 gRNA interactome also included M and S proteins, likely indicative of genomes associated with assembling virions. Interestingly, M and S proteins were not found in any of the other tier 1 interactomes but were also found in tier 2 of ORF8 sgRNA at 24-hpi. Replicase Polyprotein 1ab (Rep) was, as expected, also identified in the tier 1 protein interactomes affiliated with both the gRNA and N sgRNA pools, albeit only at 24-hpi. Rep is a polyprotein that, upon translation, auto-proteolyzes to produce 16 NSPs.^32^ While 270 unique peptides were identified as corresponding to Rep, due to the nature of bottom-up proteomics it was not possible to determine if the tryptic peptides correlated to Rep polyproteins or individual NSPs.

### Gene Ontology Enrichment Analysis of RNA-Protein Interactomes

Using gene ontology (GO) term enrichment algorithms, we next evaluated each interactome for enrichment of host cell proteins involved in specific biological processes to infer which pathways the specific RNAs may be involved with. This analysis revealed over-representation of several GO terms in the interactomes of each viral RNA as well as changes in over-represented GO terms between the two time points for single RNA interactomes (Figure 3, Supplementary Table 3). All interactomes were highly enriched for RNA binding proteins, with most interactomes specifically enriched for 3’ and 5’ UTR binding proteins consistent with full length RNA being present in each sample. Additionally, the N, S, and ORF8 sgRNA pools were all enriched for translation and translation initiation proteins. Terms related to viral translation and general “viral processes” were also enriched in the 8-hr interactome of S sgRNA and both 8- and 24-hr interactomes of N and ORF8 sgRNAs. Other categories of enriched terms were those related to the host cell response to viruses. Terms related to antiviral defense responses were enriched in the 8-hr interactome of the gRNA and the 24-hr interactomes of the gRNA and N Rand ORF8 sgRNA. One particularly interesting observation was an abundance of terms related to posttranscriptional gene silencing by RNA. All viral RNAs had at least two proteins identified that were associated with this term at both time points and four proteins related to posttranscriptional gene silencing by RNA (TRIM71, CELF1, MOV10, and LIN28B) were found in at least 3 independent RNA interactomes at any one single time point. Another highly enriched term was cytoplasmic ribonucleoprotein granules. This term, which encompasses machineries that regulate either the formation of cytoplasmic P-bodies (sites of RNA silencing) or stress granules (sites of translation suppression), was enriched in all four interactomes at either time point. Cytoplasmic ribonucleoprotein granule proteins identified in all four interactomes at both time points included APOBEC3F, MOV10, IGF2BP1, and L1RE1.

**Figure 3.**
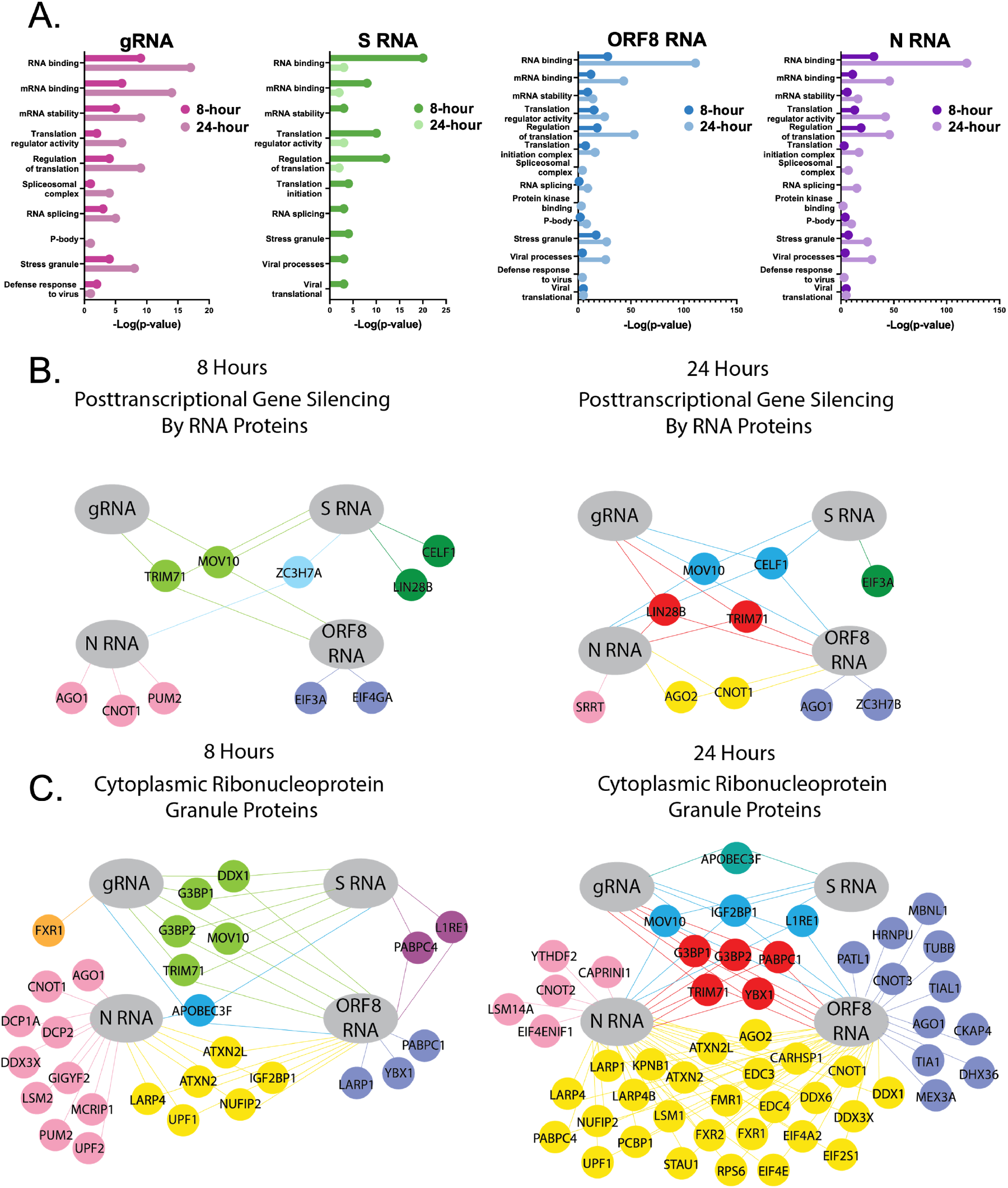
Analysis of SARS-CoV-2 RNA Protein Interactomes. (A) Gene ontology enrichment analysis of the genomic, S, N, and ORF8 RNA protein interactomes at both 8- and 24-hour time points. Refer to supplementary table 3 for complete data. (B) Proteins identified as related to “Posttranscriptional Gene Silencing” according to GO analysis using STRING plotted to show which interactomes they were identified in at the 8- and 24-hour time points. (C) Proteins identified as related to “Cytoplasmic Ribonucleoprotein Granule” according to GO analysis plotted to show which interactomes they were identified in at the 8- and 24-hour time points. Colors in (B) and (C) distinguish proteins unique to or shared among the indicated viral RNA interactomes.

We also compared the 509 proteins identified in one or more of the gRNA and sgRNA interactomes to the 617 SARS-CoV-2 RNA-associated proteins reported in six other recently published studies.^33–38^ This comparison revealed that nearly half of the proteins we identified were previously identified in at least one other study, and more than a quarter were identified in at least two (Supplementary Table 2). Additionally, GO term analysis of the proteins identified both in this study and at least one other showed enrichment of many of the same GO terms identified in Figure 3. These included mRNA stabilization and processing, translation initiation, cytoplasmic ribonucleoprotein granules, and posttranscriptional gene silencing by RNA (Supplementary Table 3). The overlap in both identified proteins as well as the enriched GO terms provided further validation for the efficacy of our HyPR-MS_SG_ approach and the potential importance of the pathways we identified. Novel protein interactors included known RNA binding proteins such as DDX20, DDX21, and YTHDF2 as well as a range of novel translation initiation factors. Of specific interest were the proteins TRIM71 and APOBEC3CF which were associated with gene silencing by RNA and ribonucleoprotein granules, respectively, both major categories of interest.

### Validation of a subset of viral RNA interactors using siRNA silencing

To determine the potential relevance of a subset of RNA interacting proteins identified using HyPR-MS_SG_ to the SARS-CoV-2 viral life cycle, five proteins predicted to be components of the host antiviral response were selected for siRNA knockdown to assess their effect on viral RNA and protein levels (Figure 4). Additionally, infectivity was measured using a reporter virus engineered to express the mNeonGreen protein. The proteins selected for follow-up analysis were chosen based on their presence in at least two independent viral RNA interactomes, as well as their GO annotations suggesting antiviral functions. Three of the proteins, MSI2, LIN28B, and PPP1CC, had previously been identified as interacting partners of SARS-CoV-2 RNA.^33,34,38^ The other two proteins selected, TRIM71 and APOBEC3F, were novel viral RNA interactors. LIN28B and TRIM71 are known to regulate posttranscriptional gene silencing by RNAs, and, along with MSI2, also regulate mRNAs.^39–41^ APOBEC3F is part of the innate immune defense targeting the RNA and DNA of several viruses.^42^ PPP1CC is from a family of protein phosphatases known to be viral transcription regulators.^43^ TRIM71, APOBEC3F, and PPP1CC are all known to play a role in cytoplasmic granule formation.^44,45^

**Figure 4.**
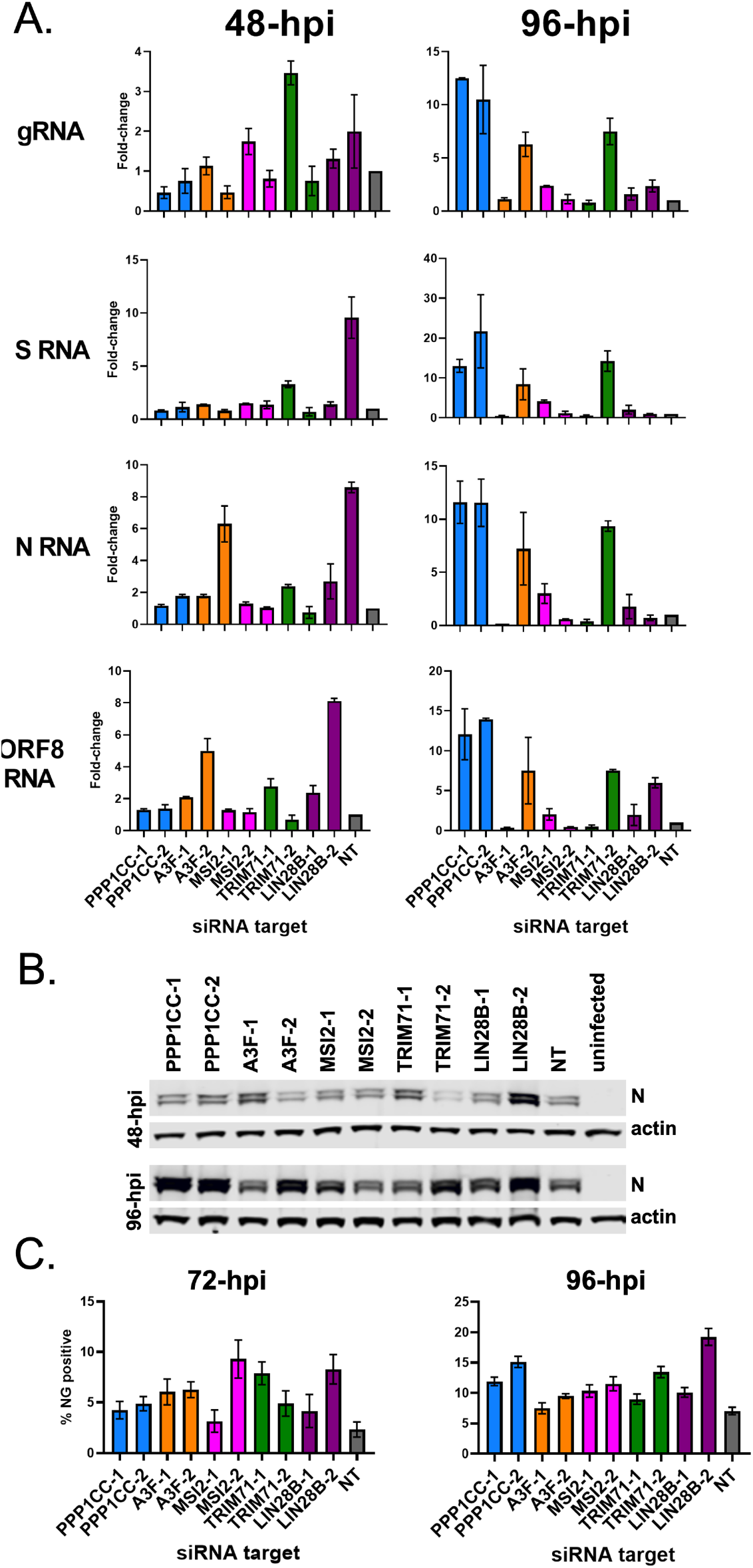
Effects of siRNA knockdowns on viral replication readouts. Calu3 cells were transfected with two independent siRNAs targeting PPP1CC, A3F, MSI2, TRIM71 and LIN28B as well as a non-targeting siRNA control. 48h post-siRNA, cells were infected with SARS-CoV-2 and harvested at the indicated time points for analysis of viral RNA expression (A), viral protein expression (B) or the percentage of NG positive cells (C). (A) RT-qPCR results showing the fold-change for each SARS-CoV-2 RNA 48- and 96-hpi normalized to 18S RNA after siRNA-mediated knockdown of the indicated targets. Fold-changes are relative to a non-targeting (NT) control set to 1. (B) Western blot analysis detecting SARS-CoV-2 N protein after siRNA-mediated knockdown of the indicated targets. (C) Flow cytometry analysis of infected cells showing the percentage of NG-positive cells at the indicated time points post-infection. For (A) and (C) data relative to a non-targeting siRNA control (set to 1) and error bars represent the standard error of the mean (SEM).

For this analysis, two siRNAs were used to target each mRNA encoding the protein of interest in the human lung epithelial cells (Calu-3) and effects on viral RNA and protein expression as well as viral spread assessed. Note that our initial screening was done in Huh-7 cells overexpressing ACE2, which substantially increases permissiveness to SARS-CoV-2 resulting in rapid viral spread. We reasoned that the Calu-3 lung cells that endogenously express ACE2 and support slower virus spread represent a more relevant model to test the contributions of the putative regulatory RNA-binding proteins to SARS-CoV-2 replication. In this model, knockdown of host proteins resulted in increased viral RNA levels for at least one of the siRNAs tested, with the largest increases observed after 96 hours for all but LIN28B (Figure 4A, Supplementary Figure 1). The PPP1CC knockdown consistently showed the largest increase, with both siRNAs resulting in more than 10-fold increases for all measured viral RNAs at the 96-hour time point. TRIM71, APOBEC3CF, and MSI2 had viral RNA levels increasing between 2- and 7-fold at 96-hpi. LIN28B had the largest changes (around 6-fold) at 48-hpi. The large increases in viral RNA levels observed suggested that the knocked-down proteins play an antagonistic role in the viral life cycle. Importantly, a similar trend was seen in western blot analysis of the SARS-CoV-2 N protein and FACS analysis of cells infected with the SARS-CoV-2 NG reporter virus, where at 96-hpi most of the siRNA knockdown conditions had resulted in increases to both N protein levels and expression of the viral mNeonGreen reporter (Figure 4B, C). The percentage of cells expressing viral NG reporter was statistically significant in at least one siRNA assay for all targets except APOEBEC3CF with statistically significant increases between 4.5% to 12.2% compared to the NT control.

## DISCUSSION

A major challenge in SARS-CoV-2 research has been differentiating between the specific interactomes of the array of RNAs the virus produces. Here, we successfully resolved the specific protein interactomes for four distinct viral RNAs at both 8- and 24-hpi. To our knowledge, this is the first description of the specific interactomes of the sgRNAs N, S, and ORF8. We identified over 500 proteins including both previously identified interactors and novel interactors. Most of the identified proteins were not identified in all interactomes, and many were uniquely identified in association with a single viral RNA-protein interactome. These unique proteins offer possible insight into the individual functions of each RNA as a part of the SARS-CoV-2 life cycle. Broadly, our findings highlight several host RNA-binding proteins that may regulate key aspects of SARS-CoV-2 replication and antiviral host defenses.

Looking to some of the pathways identified as enriched, one finding was the presence of proteins related to posttranscriptional gene silencing by RNA. Posttranscriptional gene silencing by RNA works by either destabilizing mRNAs or inhibiting translation. It encompasses silencing mediated by microRNAs (miRNAs), small interfering RNAs (siRNAs), and long noncoding RNAs (lncRNAs). RNA-interference (RNAi) is a known antiviral response by host cells that most commonly occurs when gene silencing is initiated by dicer-dependent production of virus-derived small interfering RNAs (vsiRNAs.) The vsiRNAs then target the viral RNAs in the cell for degradation.^46^ Previously, the N and ORF7A proteins of SARS-CoV-1 have been identified as suppressors of RNAi in host cells.^47,48^ TRIM71 has been shown to associate with the AGO2 protein and functionally interact with miRNAs to modulate the cell cycle.^49^ Additionally, LIN28B is known to regulate miRNA biogenesis and has also been identified as a regulatory protein that can modulate mRNA translation.^50^

Another notable observation from GO term analysis is that all RNAs interactomes are highly enriched for cytoplasmic ribonucleoprotein granule proteins. Cytoplasmic ribonucleoprotein granules form when mRNAs and proteins condense in the cytoplasm sequestering ribosomal components, translation initiation factors, and other RBPs. The two most common examples of this are processing bodies (P-bodies), which contain mRNAs that have been targeted for degradation, and stress granules, which contain mRNAs that have been translationally stalled.^51^ These bodies can form as a cellular defense against viral infection, and in at least some cases involve kinase activation.^52,53^ Alternatively, viruses can hijack stress granules to counter the cell’s defenses or even promote viral replication. At 8-hpi, gene ontology analysis showed all four interactomes were enriched for cytoplasmic ribonucleoprotein granules, and at 24-hpi, all but the S sgRNA interactome were enriched for proteins in this pathway. These included proteins thought to be nucleating factors for granules, such as G3BP1 and G3BP2, as well as additional proteins that contribute to the diversity of cytoplasmic ribonucleoprotein granule functions such as translation initiation factors. We found it notable that viral nucleocapsid protein N has been previously identified as interacting with proteins G3BP1 and G3BP2 and has been shown to disrupt stress granule formation by localizing with the proteins and sequestering them from their binding partners. Stress granule disassembly is thought to promote viral replication, but the mechanism by which this occurs is not fully understood.^54–59^ Previous studies have observed a relationship between SARS-CoV-2 RNA and proteins associated with stress granule formation. However, whether this is the result of the cell’s innate immune defense or a means of promoting viral replication remains unclear.^37,38^ P-bodies are also known to be associated with innate immune defense. When mRNAs are silenced by miRNAs, they typically localize to P-bodies for storage or degradation.^60^ Moreover, antiviral defense proteins APOBEC3G and 3F are concentrated in P-bodies and can accumulate in stress granules when cells undergo stressful events. PPP1CC is also known to translocate to stress granules and phosphorylation is an important regulator of P-bodies and stress granules.^61^ TRIM71 is also known to localize to P-bodies during its interactions with both mRNAs and miRNAs.^49^

The siRNA knockdowns of host proteins from the RNA-interactome data resulted in significant increases to the levels of all four measured viral RNAs, N protein, and infectivity. Given that the pathways we selected these proteins from were primarily stress responses or antiviral, these results were consistent with our initial prediction that the knockdowns would result in increases of viral replication measurements. However, we do point out challenges related to interpretating the siRNA knockdown results due to inconsistency in the siRNA knockdown assays. Two different siRNA assays were performed for each of the protein targets, and aside from the PPP1CC knockdown, all the assays yielded different results in magnitude and timing of the siRNA knockdown effect (Figure 4). It is unclear whether this is biologically meaningful or the result of ineffective siRNA knockdowns. We were unable to identify robust antibodies needed to confirm depletion of the targeted proteins by western blot and qPCR analysis of knockdowns was inconclusive for many of the targets. Additionally, there can be a non-linear relationship between RNA and protein expression levels further complicated analysis. That said, while not all the proteins validated were identified as interactors of all four viral RNAs, effects of the protein knockdown were uniformly observed for all viral RNAs, and at a similar magnitude, consistent with the interdependent nature of the viral life cycle. The data was further corroborated by the observed increases to the levels of the viral N protein and viral NG reporter gene expression.

Despite these limitations, interesting observations about the functional role of this subset of interacting proteins on the viral life cycle can still be made. Specifically, in APOBEC3F, PPP1CC, and TRIM71, one or both assays were able to establish statistically significant increases (p-value < 0.05) in all four of the viral RNAs at either the 72- or 96-hour time points. Both APOBEC3F and TRIM71 are novel interactors, and this data suggests not only are they binding to the viral RNA but also are functioning as viral antagonists. APOBEC3F comes from a family of viral antagonists that work by directly modifying viral DNA or RNA, specifically in RNA modifying uracil to cytosine. Genetic analysis of SARS-CoV-2 has established an overrepresentation of uracil to cytosine mutations which some have hypothesized could be related to a protein in the APOBEC family.^62,63^ Our analysis provides evidence that APOBEC3F could be one of the proteins responsible for this. The other novel RNA interactor, TRIM71, also comes from a family of known viral restriction factors. However, TRIM71 is also known to be a regulator of miRNAs and can work as a transcriptional repressor.^64^ While it is apparent that TRIM71 knockdown results in increased viral RNA expression, its varied functions mean that the mechanism by which this occurs requires further research. By far the largest effect on viral RNA levels was seen after the knockdown of PPP1C, a catalytic subunit of protein phosphatase 1 known to be involved in both HIV and Ebola virus transcription.^65,66^ Future research on these three proteins as well as MSI2 and LIN28B (which also showed statistically significant changes in viral RNA levels) should include work to localize the RNA-protein interactions as well as additional functional assays to more specifically determine how protein knockdowns affect other measures of viral fitness.

Overall, this study describes and validates a powerful strategy to exploit the unique leader junction of SARS-CoV-2 sgRNAs to purify them with specificity and characterize their protein interactomes. We also provide an unprecedented description of the protein interactomes of N, S, and ORF8 sgRNA and full-length gRNA across two time points, representing early and late infection. Analysis of the identified proteins provided insight into several biological pathways SARS-CoV-2 interacts with throughout the course of infection, specifically in the case of posttranscriptional gene silencing by RNA and cytoplasmic ribonucleoprotein granules. Using siRNA knockdown assays, the effect of knocking out five host proteins (including two that had not previously been identified as SARS-CoV-2 interactors) was observed at the levels of viral RNA abundance, N protein expression, and net infectivity. Statistically significant increases of viral RNA levels were observed across all siRNA knockdown of host protein targets, showing the potential functional importance of these interacting partners to the viral life cycle. This study provides significant insight into the host protein-viral RNA interactome and particularly the role of each individual RNA throughout the course of SARS-CoV-2 infection.

## Supporting information

Supplementary Table 1 presents all qPCR assay sequences, siRNA assay information and capture/release oligonucleotide sequences.

Supplementary Table 3 presents gene ontology analysis results.

Supplementary Table 4 presents raw qPCR values from RNA capture experiments.

Supplementary Table 5 presents raw FACS values from siRNA knockdown analysis and ANOVA test results.

Supplementary Table 6 presents raw qPCR values from siRNA knockdown analysis and ANOVA test results.

## DATA AVAILABILITY

The mass spectrometry proteomics data have been deposited to the ProteomeXchange Consortium via the PRIDE partner repository with the dataset identifier PXD040147.^67^

## SUPPLEMENTARY DATA

Supplementary Table 1 presents all qPCR assay sequences, siRNA assay information and capture/release oligonucleotide sequences. Supplementary Table 2 presents proteomics results from Student’s t-tests and analysis of other SARS RNA-protein interactomes studies. Supplementary Table 3 presents gene ontology analysis results. Supplementary Table 4 presents raw qPCR values from RNA capture experiments. Supplementary Table 5 presents raw FACS values from siRNA knockdown analysis and ANOVA test results. Supplementary Table 6 presents raw qPCR values from siRNA knockdown analysis and ANOVA test results. Supplementary Figure 1 shows complete qPCR analysis from all four siRNA knockdown time points.

## TABLE AND FIGURES

**Supplementary Figure 1. Effects of siRNA knockdowns on viral RNA levels**. Calu3 cells were transfected with two independent siRNAs targeting PPP1CC, A3F, MSI2, TRIM71 and LIN28B as well as a non-targeting siRNA control. RT-qPCR results showing the fold-change for each SARS-CoV-2 RNA at 24-, 48-, 72-, and 96-hpi normalized to 18S RNA after siRNA-mediated knockdown of the indicated targets. Fold-changes are relative to a non-targeting (NT) control set to 1. Error bars represent the standard error of the mean (SEM).

